# No single measure is enough: Recovery of the Critically Endangered *Mobula mobular* requires integrated maximum bycatch mitigation and nursery area protection

**DOI:** 10.64898/2026.08.18.744841

**Authors:** Mayuri Chopra, Roberto Salguero-Gómez, Guy M. W. Stevens, Gwilym Rowlands, Divya Karnad, T. Mohanraj, Daniel Fernando, Katrina J. Davis

## Abstract

As anthropogenic threats have intensified over the past 500 years, we find ourselves in the midst of a sixth mass extinction, with continued losses of biodiversity threatening ecosystem stability. This biodiversity loss has caused species extinctions across taxa, and placed several others at high risk of functional extinction. These disturbance-driven impacts represent one of the most acute biodiversity crises facing global marine systems. Species exhibiting slow life histories characteristically have low resilience to disturbance. Here, we assess the risk of functional extinction and identify policy pathways for population recovery of the slow-living, Critically Endangered elasmobranch, the spinetail devil ray (*Mobula mobular*). We develop a stochastic, state-structured Integral Projection Model (IPM) parameterised with demographic data collected from fishery landings data in India, the world’s largest mobulid fishery, and supplemented with data on vital rates from published literature. Using the IPM, we estimate that the population is declining at approximately 12% annually, experiencing substantial limiting pressure from fisheries overexploitation and failing to approach its biological maximum growth potential. Our results indicate that populations of *M. mobular* will be at high risk of functional extinction if ‘business as usual’ harvest scenario persists for another decade. We further show that long-term population recovery is only possible if survival increases significantly across all size classes, especially among large reproductive females, alongside a concurrent increase in fecundity. We conclude that no single policy measure is sufficient to recover population of *M. mobular* along the southeastern coast of India. Instead, combined protection through maximum bycatch mitigation and protection of nursery areas in no-take zones will be required for population recovery. This research demonstrates that recovery of overexploited populations often requires integrated resource management across life stages, and that the Critically Endangered *M. mobular* warrants urgent conservation action to avoid functional extinction.

## 3. Introduction

Overexploitation of marine resources drives global biodiversity loss (Crutzen, 2002; Jungblut et al., 2020). This loss threatens ecosystem stability and increases risk of species extirpation (Barnosky et al., 2011; Odia, 2025). Intensification of anthropogenic threats over the past 500 years has led to >1,358 marine species being listed as Vulnerable, Endangered, or Critically Endangered on the International Union for the Conservation of Nature (IUCN) Red List of Threatened Species (Lotze, 2021). Disturbances driving marine biodiversity loss include habitat degradation, pests and disease, climate change, and overharvesting of stocks (Ward et al., 2022). These disturbance-driven impacts represent one of the most acute biodiversity crises facing global marine systems (O’Hara et al., 2021).

Resilience to human disturbances is particularly low for taxa with slow life histories. Slow life histories are characterised by large size, high adult survival, low fecundity, and slow developmental rates (Bennett and Owens, 1997; Stearns, 1998; Sæther and Bakke, 2000; Healy et al., 2019). Examples include the white shark (*Carcharodon carcharias*), which can live up to 70 years (Hamady et al., 2014), and the Greenland shark (*Somniosus microcephalus*), which may live up to 400 years (Nielsen et al., 2016). A key feature of slow life histories is the “malediction” (*sensu* Lebreton, 2006): an inverse relationship between generation time and maximum population growth rate, often driving populations into a so-called extinction vortex (Gilpin and Soule, 1986). An extinction vortex is a progressive decline in population viability driven by interacting environmental, genetic, and demographic factors that create a feedback loop toward extinction (Gilpin and Soule, 1986; Brook et al., 2008). Intense disturbances, such as overfishing, can decrease population growth, increasing vulnerability to subsequent disturbances and potentially inducing extinction vortices (*sensu* Lacy and Lindenmayer, 1995) in slow-living species (Gamelon et al., 2014). Disturbance impacts on slow-living species’ recovery potential can propagate across generations (Lotze et al., 2011), with populations exhibiting elevated extinction risk (Dulvy et al., 2003) or irreversible collapse (Keevil et al., 2018). Beyond biological extinction, slow-living species are at risk of functional extinction (Shaffer, 1981; Dulvy et al., 2003; Jarić et al., 2016). Here, functional extinction follows Shaffer’s (1981) minimum viable population (MVP) concept, below which demographic, environmental, and genetic stochasticity may prevent population persistence. Thus, understanding how disturbances affect the demographic processes across a species’ life cycle, which define its life history strategy, is critical for assessing recovery potential and informing conservation strategies for slow-living species (Pirotta et al., 2018; Morris et al., 2006).

Demography of slow-living shark and ray taxa has been widely studied to address their urgent conservation concerns (Mejía et al., 2025). Population models have been extensively used to examine elasmobranch responses to anthropogenic stressors, such as overfishing (Stevens et al., 2000; Frisk et al., 2005; Dulvy et al., 2014; Pacoureau et al. 2021). Examples include correlating extinction risk with population growth rates in sharks and rays (Simpfendorfer, 2005; Mejía et al., 2025), estimating rebound potential for Atlantic sharpnose shark (*Rhizoprionodon terraenovae*), porbeagle (*Lamna nasus*), and Pacific spiny dogfish (*Squalus suckleyi*) (Au et al., 2015), and large coastal and pelagic shark species: blacktip shark (*Carcharhinus limbatus*) and silky shark (*Carcharhinus falciformis*) (Crouse et al., 1987; Cortés, 1998; Grant et al., 2020). Demographic parameters from these models, such as population growth rate (*λ*), are fundamental for evaluating species resilience to overexploitation (Pardo et al., 2016; de Barros et al., 2022). Demographic studies of sharks and rays have predominantly relied on matrix population models, which structure populations using discrete life-cycle stages (Caswell, 2001; Cortés, 2002; Simpfendorfer et al., 2005; Gedamke et al., 2007; Coelho et al., 2015; Smallegange et al., 2016). While these models are widely applied and efficient for species truly structured into discrete life-cycle stages, such as cheloniid sea turtles (egg, hatchling, juvenile, adult) (Crouse et al., 1987), they artificially discretise otherwise continuous traits (such as size) (Merow et al., 2014).

Integral Projection Models (hereafter IPMs; Easterling et al., 2000) are demographic models that examine population dynamics in discrete time and continuous traits (*e.g*., size). Since 2000, IPMs have gained popularity due to their flexibility in incorporating discrete and continuous traits (Merow et al., 2014), efficiency when applied to small demographic datasets (Ramula et al., 2009), and usefulness in testing ecological hypotheses (Coulson, 2012). To date, IPMs have been applied to over 300 species (Levin et al., 2022). Despite their advantages, IPMs have rarely been applied to shark and ray conservation, although these species’ life cycles are best represented using both discrete stages and continuous traits (for example, age and reproductive stages, as in Hussey et al., 2010). Global accounts of IPMs applied to fisheries management of endangered elasmobranchs are sparse, with only three accounts of using Dynamic Energy Budget Integral Projection Models (DEB-IPMs) on elasmobranchs (Smallegange et al. 2016, Smallegange et al. 2020, Lucas et al., 2025).

Here, we assess the risk of functional extinction and identify policies for population recovery of the slow-living elasmobranch; the spinetail devil ray (*Mobula mobular*). The monogeneric family Mobulidae, are recognised as the ray family most vulnerable to fishing pressure (Mejía et al., 2025), particularly in the Indian Ocean (Barrowclift et al., 2025). Population declines of up to 99% (CITES, 2025; Laglbauer et al., 2026) have prompted an international trade ban under the Convention on International Trade in Endangered Species of Wild Fauna and Flora (CITES (2025), and uplisting to Critically Endangered on the IUCN Red List of Threatened Species (Jabado et al., 2025a, 2025b, 2025c). We first develop a stochastic state-structured IPM parameterised with demographic data from fishery landings in India, the world’s largest mobulid fishery (Laglbauer et al., 2026), and vital rates from the literature. Using this IPM, we then test the following three hypotheses: (H1) Populations of *M. mobular* will be at high-risk of becoming functionally extinct if current fishing pressure persists for another decade. This hypothesis is motivated by the on-going population decline of 86% over the past decade (Chopra et al., 2026), indicating that further declines of similar magnitude have a high risk of compromising population viability due to reduced resilience of slow-living species following disturbances (Gamelon et al., 2014); (H2) Existing conservation measures are inadequate to facilitate population recovery unless they substantially increase adult survival for multiple generations, as recovery in highly exploited stocks of slow-living species is slow and typically expected only over multiple generation times (Flowers et al., 2020); and (H3) Populations of *M. mobular* exhibit high sensitivity to adult survival and have a heightened functional extinction risk when adult survival is low. This expectation is motivated by the empirically observed higher sensitivity of slow-living species to changes in adult survival (Heppell et al., 1999; Kopf et al., 2025).

## 4. Materials and methods

### 4.1 Data collection

#### 4.1.1 Study site

Part of the demographic data utilised in this study comes from a high–bycatch-rate fishery in the southeastern coast of India (8.8165° N, 78.1625° E and 8.8922° N, 78.1707° E) (Supplementary Information 1.1; Fig. 1).

**Figure 1.**
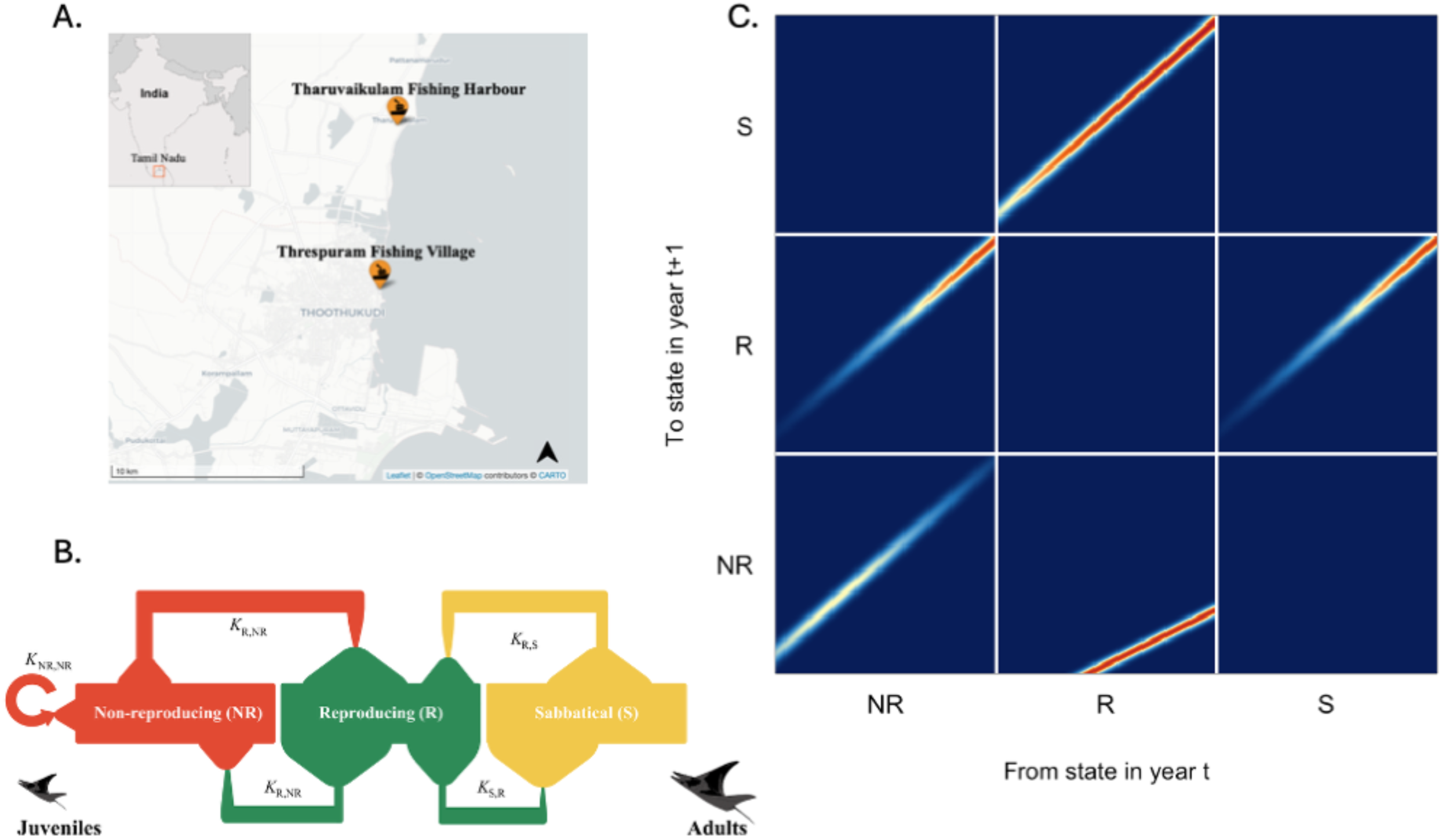
Stochastic state-structured Integral Projection Model (IPM) parameterised for *Mobula mobular* female population. **A.** Fish landing centres in Tamil Nadu, in the southeastern coast of India. **B.** The life cycle of the female populations of *M. mobular*, used to parameterise our state-structured IPM kernel. Red, green and yellow indicate transition between individuals in reproductive states, Non-reproductive (NR), Reproductive (R) and Sabbatical (S), respectively. **C.** The state-structured IPM kernel with a 3×3 Goodman matrix consisting of kernels with transitions between states: Non-reproductive (NR) state constitutes individuals which have not started reproducing yet; Reproductive (R) are those producing pups; and Sabbatical (S) constitutes females resting between two pregnancies. The colour scale ranges from deep blue (lowest values) through yellow (mid-range values) to red (highest values). Each kernel represents the survival, size-change and transition between two states of the population, within the size ranges estimated in disc width (cm). The kernels in the block matrix are denoted as: (1) K_NR,NR_ (Non-reproductive remaining as Non-reproductive), (2) K_R,NR_ (Non-reproductive to Reproductive) (3) K_S,NR_ (transition does not exist), (4) K_NR,R_ (Reproductive individuals producing Non-reproductive pups), (5) K_R,R_ (transition does not exist), (6) K_S,R_ (Reproductive to Sabbatical), (7) K_NR,S_ (transition does not exist), (8) K_R,S_ (Sabbatical to Reproductive), (9) K_S,S_ (transition does not exist).

#### 4.1.2 Demographic data

To parameterise the IPM for populations of *Mobula mobular*, we used empirically collected field data, vital rates published in the literature for *M. mobular*, and vital rates from closely related species within the *Mobula* genus when species-specific rates were unavailable (Supplementary Information 1.2). The spinetail devil ray dominates 86% of the mobulid fishery in the region (Chopra et al., 2026) and shares a similarly slow life history with other overexploited mobulids (Barrowclift et al., 2025). *Mobula mobular* individuals exhibit slow-living through their late maturity (6–10 years) (Pardo et al., 2016; Barrowclift et al., 2025), a long gestation period (approximately annual) (Kiyatake et al., 2025), low reproductive output (pup biennially or triennially) (Rambahiniarison et al., 2018), and moderately long life spans (20–25 years) (Barrowclift et al., 2025). As such, we assumed the populations of *M. mobular* are representative of the relative risk across the *Mobula* genus. We parameterised the IPM for *M. mobular* females as is common practice in animal demography for species with separated sexes (Rees et al., 2014; Macdonald et al., 2023).

### 4.2 Model construction

To test our hypotheses regarding (H1) populations of *M. mobular* will be at high-risk of becoming functionally extinct if current fishing pressure persists for another decade, (H2) existing conservation measures are inadequate to facilitate population recovery unless they substantially increase adult survival for multiple generations, and (H3) populations of *M. mobular* exhibit high sensitivity to adult survival and have a heightened functional extinction risk when adult survival is low, we constructed a stochastic state-structured IPM (Easterling et al., 2000) for *M. mobular*. Briefly, a standard IPM examines population dynamics in discrete time (*e.g.*, between *t* and *t+1*), where vital rates of survival, growth, and reproduction are functions of a continuous trait such as size. Given the complex life cycle of *M. mobular*, we developed a model with three discrete states linked to reproduction (‘Non-reproductive’ [NR], ‘Reproductive’ [R], and ‘Sabbatical’ [S]), where each individual within each discrete class is also quantified by its size, in this case disc width (Fig. 1). The three-state structure of our IPM allowed us to clearly separate non-reproductive from reproductive individuals to explicitly examine the delay periods between maturity and reproduction (Rambahiniarison et al., 2018), which is crucial to test population response to harvest and recovery scenarios (H1 and H2). As previously highlighted, individuals show biennial or triennial reproduction, taking a full year of sabbatical after reproducing (Pardo et al., 2016; Rambahiniarison et al., 2018). Consequently, the Non-reproductive (NR) state constitutes juveniles as well as mature females who can become pregnant but have not yet reproduced, and the Reproductive (R) state constitutes females that are pregnant and produce a pup in year *t*+1. Finally, the Sabbatical (S) state consists of females that are resting after having produced a pup in year *t*, but will return to the Reproductive (R) state after the annual break period, if they survive (Fig. 1).

#### 4.2.1 Vital rate estimation

To build the IPM, we first estimated parameters constituting the vital rates of survival, growth, and reproduction for the lifecycle of *M. mobular* (Fig. 1) (Supplementary Information 1.3).

#### 4.2.2 Integral Projection Model building and matrix assembly

We used the estimated annual vital rate parameters and modelled functions in section 4.2.1 (Supplementary Information 1.3; Table 2) to construct a 3×3 Goodman matrix (Goodman, 1969) format to accommodate the IPM kernels for survival-dependent growth (*P* sub-kernel) and reproduction (*F*) associated with each of the three states: Non-reproductive (NR), Reproductive (R), and Sabbatical (S). The *P* sub-kernel within this 3-discrete state IPM consists of the probability of survival (*σ*(*z*)) and growth (*γ*(*z*)), such that:

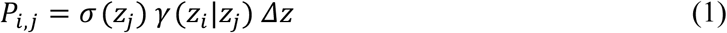

Where *P_i,j_* is the transition probability from size *z_j_* at time *t* to size *z_i_* at time *t+1* conditional on having survived between *t* and *t*+1, *σ*(*z_j_*) is the survival probability, *γ* (*z_i_* | *z_j_*) is the growth probability density of *z_i_* at *t+1 given* size *z_j_* at *t*, and *dz* is the size-class width.

The *F* sub-kernel of the IPM describes the recruitment of new female pups of a given size in *t*+1 by a mother of size *z_j_* at time *t.* Here, recruit sizes were drawn from a Gaussian distribution with mean *μ_p_(z_i_)* and variance *σ*^2^*_pup_*, scaled by a constant probability of successful recruitment. The reproduction probability (*φ*(*z*)) was implicit when an individual transitions between the NR, R and S states, such that:

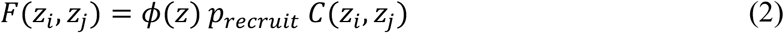

where the offspring size distribution is,

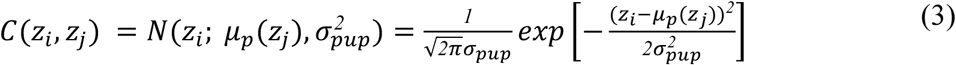

Where *z_j_* is mother size at time *t* and *z_i_* is offspring size at *t+1* and *p_recruit_* is the probability that a pup survived and was recruited in the population. We modeled offspring size as a function of mother size because mobulids have matrotrophic reproduction, meaning that mothers directly nourish the embryo, and nutrition for the embryo is not solely derived from the yolk sac (Blackburn and Hughes, 2024). As such, larger mothers, which naturally have larger maternal standard length and body fat have larger offspring size at birth (Cortés, 2000; Hagmayer et al., 2018).

Each of the nine IPM kernels accommodated in the 3×3 Goodman matrix (Fig. 1) can be expressed as per Eq. (4), following Levin et al. (2021):

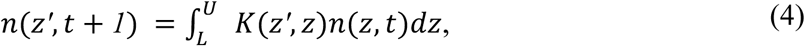

Where *n(z’, t + 1)* is the size distribution (mobulid disc width) of the population at time *t + 1*, with *z’* being the state variable used to characterise the population. The integral of Eq. (4) over the range of *z’* gives the total population size in that state (NR, R, or S), as well as its population structure. The lower and upper bounds of the integral were represented by *L* and *U*, respectively, and here were determined based on the minimum (102 cm) and maximum size (242 cm) of observed individuals landed in our census of 2023.

To incorporate parameter uncertainty in the model, we introduced annual stochasticity in all vital rate parameters including survival (*σ*_0_, *σ*_1_), growth (*L_max_*, *k*, and *γ*_(*SD*)_), reproduction probability from NR individuals (*φ*_0_, *φ*_1_), recruit size distribution (mean and standard deviation), and a size-independent vital rate constant of pup to recruitment probability. The IPM was built using 200 mesh points (for each of the three states, NR, R, and S) over the domain [*L, U*] defined in Eq. (4). The constructed IPMs were ordered in a block matrix for transitions between all states, Non-reproductive (NR), Reproductive (R), and Sabbatical (S). The block matrix consists of (Fig. 1): (1) K_NR,NR_ (Non-reproductive individuals that survive, may grow in size, but remain as Non-reproductive), (2) K_R,NR_ (Non-reproductive individuals that survive, may grow, and then transition to the Reproductive state by becoming pregnant), (3) K_S,NR_ (transition does not exist, as an individual must reproduce to take a Sabbatical), (4) K_NR,R_ (Reproductive individuals that produce pups, which are themselves Non-reproductive), (5) K_R,R_ (transition does not exist, as a pregnant *M. mobular* individual has an annual gestation period, and on survival will transition to Sabbatical), (6) K_S,R_ (Reproductive individuals that survive, may grow, and then transition to Sabbatical), (7) K_NR,S_ (transition does not exist, as Sabbatical individuals don’t reproduce in the same year), (8) K_R,S_ (Sabbatical individuals that survive, may grow, and then become pregnant, thus transitioning to the Reproductive state), (9) K_S,S_ (transition does not exist, as Sabbatical individuals move back to the Reproductive state after their annual break).

#### 4.2.3 Initialising the population vector

The initial-conditions population vector (***n****_0_*) describes the distribution of individuals in time *t*; we needed ***n****_0_* for forecasting population projections to test (H1) and (H2). The ***n****_0_* was derived from the 2023 landings data (Supplementary Information 1.4) and is a concatenated vector comprising population vectors for each of our three states: Non-reproductive (NR), Reproductive (R), and Sabbatical (S). All juveniles were assigned to the NR state. Individuals with disc widths ≥217.8 cm, corresponding to female size at maturity (Rambahiniarison et al., 2018), were classified as adults. Since at least 50% of mature female mobulids exhibit delayed pregnancy after maturity (Rambahiniarison et al., 2018), we reassigned 50% of adults in each size bin from the NR state to the R state, while the S state was initialised as empty, as it was expected to be filled by females from the R state after reproducing and surviving. The total initial female population size (***N*_0_**) was derived by scaling a total population estimate of 1,736 individuals (standardised by sampling effort; Chopra et al., 2026) by the observed proportion of females (0.488) in 2023. The estimated ***N***_0_ based on the bycatch data was 847 female individuals.

#### 4.2.4 Stochastic population projection

To project populations to *t+1*, we introduced stochasticity in all vital rates, each perturbed with small random effects in an annual parameter draw. Each year, ***n****_0_* was multiplied by the projection matrix to produce the population in the next year. Repeating this step annually projected the population forward. We forecast the population trajectory over a defined time horizon (10 years for H1; 50 years for H2), running 400 stochastic simulations for each forecast to ensure reproducible random draws.

### 4.3 IPM analysis

#### 4.3.1 Estimating minimal viable population size threshold

To test (H1), that the *M. mobular* population will be at high-risk functional extinction if current fishing pressure persists for another decade, we estimated a minimum viable population threshold, obtaining an MVP (***N***_f_) of 2,400 females (Supplementary Information 1.5). Minimum viable population (MVP) thresholds (*sensu* Shaffer, 1981) have been widely used in conservation science to assess population extinction risk and the ability of natural populations to maintain evolutionary potential (defined as enough genetic variation for future adaptive change) (Traill et al., 2010; Clements et al., 2011; Jamieson and Allendorf, 2012; Pérez-Pereira et al., 2022). Briefly, the census population size (***N***_c_) is an estimated MVP derived from the genetic effective population size following the 50/500 rule (Franklin, 1980), whereby the genetic ***N***_e_ (a measure of a population’s genetic behaviour) should not fall below 50 in the short term or 500 in the long term. The ***N***_e_ of 500 corresponds to a total population size ***N***_c_ of about 5,000 individuals (Traill et al., 2010), empirically supported by MVP-size estimates across taxonomic groups (Traill et al., 2007; 2010). To assess *M. mobular* recovery potential across scenarios, we evaluated whether the population size approached the MVP threshold.

#### 4.3.2 Population viability under fisheries pressure

To test (H1) whether the *M. mobular* population will be at high-risk of becoming functionally extinct if current fishing pressure persists for another decade, we projected a time horizon forecast of ten years. We applied a harvest pressure scenario of magnitude similar to the last decade (Chopra et al., 2026), causing an 86% decline in the population size over a ten-year period. Following Griffith (2017), we applied this perturbation (an annual multiplier) to the vital rate functions within the *P* and *F* sub-kernels across all three states (Non-Reproductive, Reproductive, Sabbatical), independent of size, assuming non-selective bycatch of mobulids.

To test the robustness of H1 to variation in the initial population size (***N***₀), we generated harvest scenario estimates across a range of initial population sizes. This was necessary because our ***N***₀ estimate (847 individuals) was derived from bycatch data in the absence of a formal *M. mobular* stock assessment (ICAR-CMFRI, 2022). Therefore, we applied the harvest scenario (H1) over three bycatch cases modifying the initial population sizes: low bycatch case (the recorded female bycatch was 25% of the total population [***N***_0,low_ = 3,389]); medium bycatch case (the recorded female bycatch was 50% of the total population [***N***_0,med_ = 1,694]); high bycatch case (the recorded female bycatch was 75% of the total population [***N***_0,high_ = 1,130]).

#### 4.3.3 Conservation management scenarios

To assess (H2) whether existing conservation measures are inadequate to facilitate population recovery unless they substantially increase adult survival for multiple generations, we used the parameterised IPM and projected a 50-year time horizon (approximately 25 years per generation; Barrowclift et al., 2025). We incorporated perturbations to survival and fecundity (Table 1) and evaluated how alternative policies alter the contribution of vital rates to population growth rate (*λ*). To assess recovery potential, we applied six scenarios linked to management actions (Scenario A—F; Table 1), equating to partial or complete protection measures such as size-based capture limits, temporary seasonal closures, spatial protection, or a blanket ban on mobulid catch, among others (Table 1). The levels of impact (50–80%) within each scenario represented measures of partial and substantive fishery impact reduction (Griffiths and Lezama-Ochoa, 2021). We incorporated perturbations (an annual multiplier) to the vital rates within *P* and *F* sub-kernels (Griffith, 2017), and all probabilities were constrained to an upper bound of to produce biologically realistic probability estimates. These population projections identify the scenarios which prevent the risk of functional extinction and shift the population trajectory towards recovery (>***N***_f_; MVP threshold).

**Table 1.**
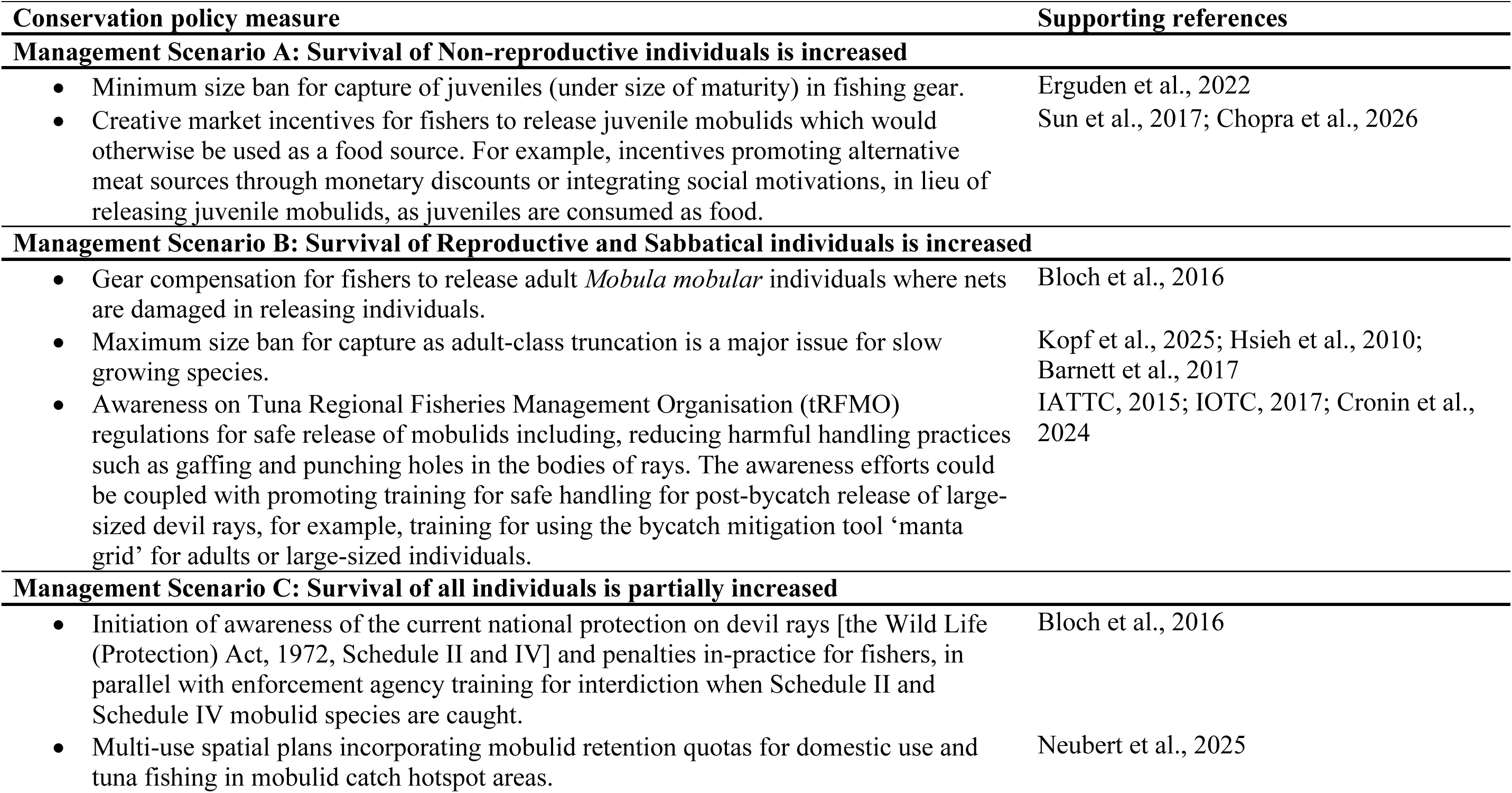
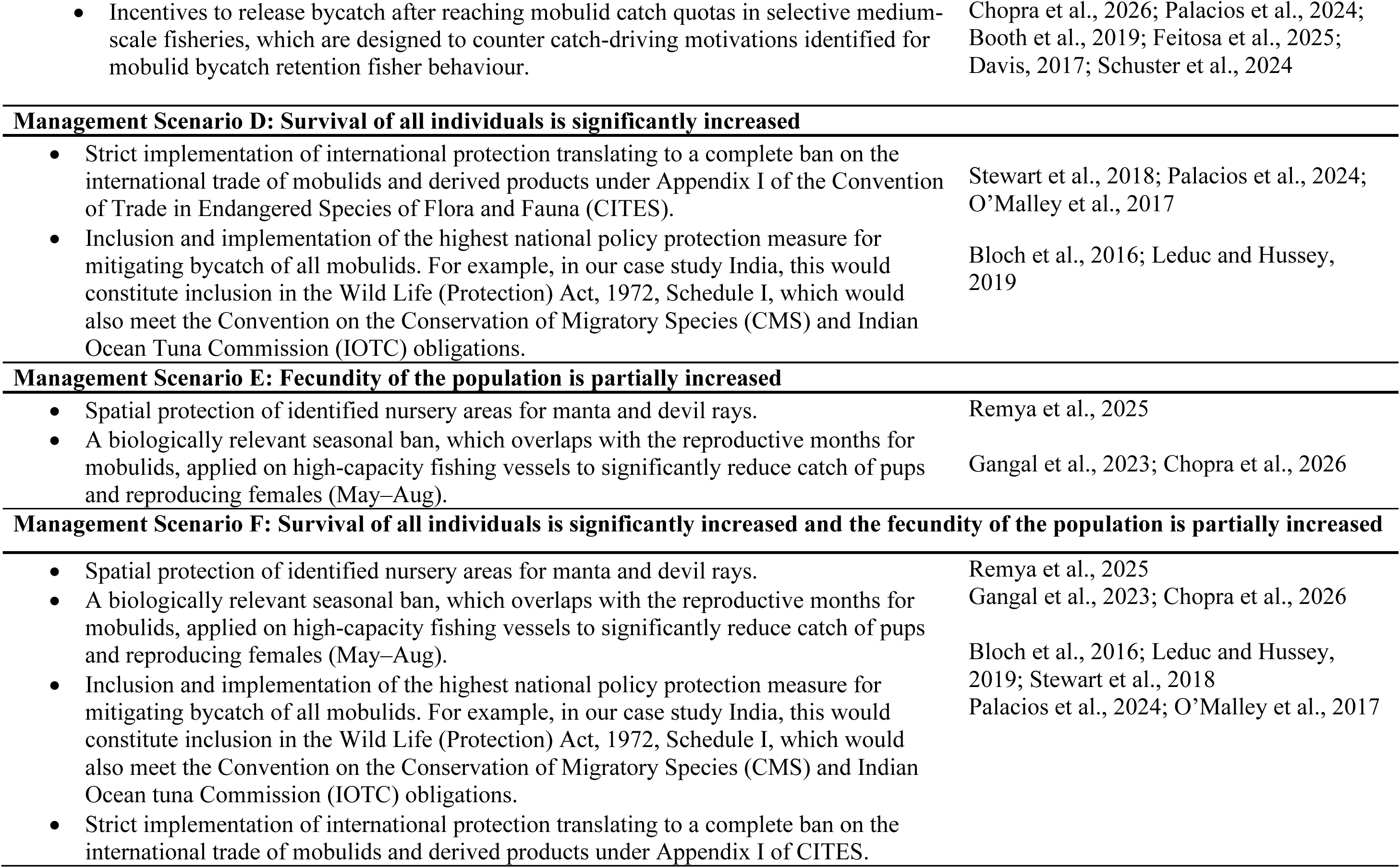
Conservation management scenarios tested with the stochastic state-structured Integral Projection Model parameterised for *Mobula mobular* populations in India. Scenario reflect policies evaluated for effectiveness in driving population recovery, aiming to identify demographically impactful conservation strategies. All scenarios were tested through stochastic projection over a 50-year time horizon.

#### 4.3.4 Sensitivity analyses

To test (H3) the populations of *M. mobular* exhibit high sensitivity to adult survival and have a heightened functional extinction risk when adult survival is low, we perturbed vital rates and assessed the effect on population growth rate (*λ*). Specifically, we calculated parameter-level sensitivities of *λ* following Griffith (2017). For each of the 19 baseline vital rate parameters for survival, growth, and reproduction (Table 1) across the three life-cycle states (Non-Reproductive, Reproductive, Sabbatical), we increased the baseline parameter value by a small additive increment (Δ*θ* = 0.0001) while maintaining all other parameters at baseline. After each perturbation, we rebuilt the full demographic model and recalculated its *λ* as the dominant eigenvalue of the discretised kernel (Ellner et al., 2016). Sensitivity of *λ* to vital rate parameter *θᵢ* was estimated as:

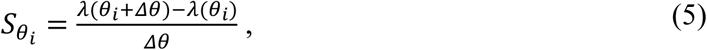

This expression quantifies the change in *λ* resulting from a small change in that vital rate parameter (*e.g*., intercept of the survival function for NR individuals). To quantify uncertainty around these sensitivity estimates, we implemented a simulation-based uncertainty propagation approach to produce a distribution of sensitivity estimates with mean and 95% uncertainty intervals (Supplementary Information 2.1).

## 5. Results

### 5.1 Stochastic state-structured Integral Projection Model

We present baseline parameter estimates used to construct the base IPM in Table 2. The base IPM, parameterised through empirical data and vital rate estimates in literature, resulted in a population growth rate (*λ*) of 0.887 (0.859, 0.881, 95% C.I.) (Table 3).

**Table 2.** Vital rate parameters, models, and estimates for the stochastic state-structured Integral Projection Model (IPM) for *Mobula mobular* populations in India. The IPM is parameterised for a female *M. mobular* life cycle, constituting its three states: Non-reproductive (NR), Reproductive (R), and Sabbatical (S). Parameters for survival, growth, and reproduction are presented with their modelled relationships, including size-dependent and size-independent parameter estimates.

| Vital rate/<br>parameter | Selected model | Parameter | Parameterisation<br>details |
| --- | --- | --- | --- |
| Survival | $\sigma(z) = \frac{l}{l + \exp[-(\sigma_0 + \sigma_1 z)]}$ | $\sigma_{0,NR} = -1.840$<br>$\sigma_{1,NR} = 0.021$<br>$\sigma_{0,R/S} = -1.630$<br>$\sigma_{1,R/S} = 0.021$ | Logistic regression (estimated using empirical data and inferred literature survival estimates) |
| Growth | $\gamma(z) = L_{\infty, \text{baseline}} - (L_{\infty, \text{baseline}} - z_t) \exp(-kdt)$ | Asymptotic size<br>($L_{\infty, \text{baseline}}$) = 245.400 cm<br>Growth coefficient ($k$) = 0.120<br>$\gamma_{(SD)} = 6.000$ | Von Bertalanffy Growth function. Asymptotic size estimated using Froes and Binohlan (2000) approach; Growth coefficient inferred from Pardo et al., 2016; standard deviation (SD) estimated from 2023 dataset. |
| Reproduction probability | $\phi(z) = \frac{l}{l + \exp[-(\phi_0 + \phi_1 z)]}$ | $\phi_0 = -6.000$<br>$\phi_1 = 0.030$ | Logistic regression |
| Recruit size | $C(z_k, z_l) = N(z_k; \mu_p(z_l), \sigma_{pup}^2)$ | $\sigma_{pup}^2 = 3.830$ | Gaussian distribution |
| Recruit to maternal size | $D_{i,t}^{pup} = 0.5D_{i,t}$ | Varies with maternal disc width (D) | Inferred from Rambahiniarison et al., 2018. |
| Pup to recruit probability | Constant | $p_{recruit} = 0.600$ | Inferred from Kashiwagi, 2014. |

**Table 3.** Estimated changes in population growth rate (*λ*) showing the highest *λ* (>1) under Scenario F. Population growth rate estimates (95% CI in brackets) are based on the stochastic, state-structured Integral Projection Model (IPM) for *Mobula mobular* population, parameterised with empirical data from India, and vital rate estimates in literature. Changes are given for the base IPM, the harvest pressure scenario, and management scenarios (A-F).

| Policy scenario | Description | Population growth rate ($\lambda$) |
| --- | --- | --- |
| Base Integral Projection Model | The base Integral Projection Model parameterised through empirical data and vital rate estimates in literature. | 0.887<br>(0.859,0.881) |
| Harvest pressure scenario | Continued harvest pressure of magnitude similar to the last decade, causing an 86% decline over 10 years. | 0.732<br>(0.697, 0.744) |
| Management Scenario A: Survival of Non-reproductive individuals is increased | 80% increase in the survival of *Mobula mobular* females in the Non-reproductive state. | 0.948<br>(0.938, 0.949) |
| Management Scenario B: Survival of Reproductive and Sabbatical individuals is increased | 80% increase in the survival of *Mobula mobular* females in the Reproductive and Sabbatical states. | 0.917<br>(0.892, 0.918) |
| Management Scenario C: Survival of all individuals is partially increased | 50% increase in the survival of *Mobula mobular* females in all three states (NR, R, and S). | 0.971<br>(0.959, 0.975) |
| Management Scenario D: Survival of all individuals is significantly increased | 80% increase in the survival of *Mobula mobular* females in all three states. | 0.981<br>(0.971,0.985) |
| Management Scenario E: Fecundity of the population is partially increased | 50% increase in the fecundity of the population. | 0.895<br>(0.872,0.893) |
| Management Scenario F: Survival of all individuals is significantly increased and the fecundity of the population is partially increased | 80% increase in survival of all individuals in addition to a 50% increase in the fecundity of the population. | 1.012<br>(1.004, 1.017) |

### 5.2 Population viability under fisheries pressure

We found support for H1: *M. mobular* population would be at high risk of functional extinction if current fishing pressure persists for another decade. Harvest pressure scenario reduced the annual population growth rate (*λ*) to 0.732 (0.697, 0.744, 95% C.I.), a 17.5% annual decrease relative to the base model (Table 3). When projected stochastically over a ten-year time horizon, populations declined sharply, with only a few hundred individuals remaining after a decade of sustained pressure equivalent to last decade’s magnitude (Fig. 2). Importantly, this sharp decline held irrespective of the initial population size (***N***₀) under low, medium, or high bycatch scenarios (Fig. 2).

**Figure 2.**
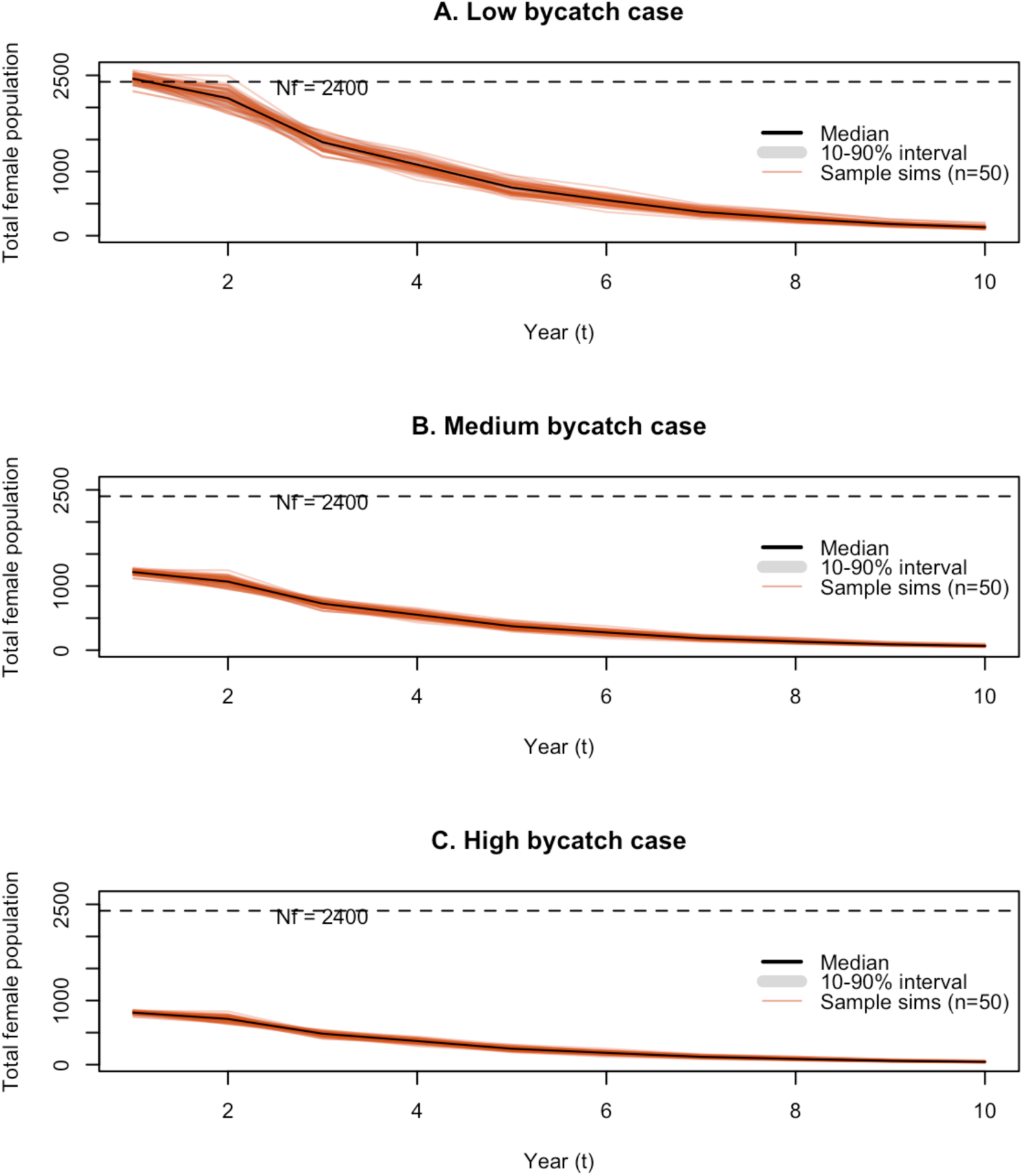
Population decline and high-risk of functional extinction under persistent fisheries pressure on *Mobula mobular* populations for the parameterised stochastic Integral Projection Model, in India. Populations are projected stochastically over 10 years of sustained fishing pressure, showing the effect of current harvest pressure under varying initial population size. **A.** low bycatch scenario, where the 2023 bycatch represents 25% of the total M. mobular population; **B**. medium bycatch scenario (50%); **C**. high bycatch scenario (75%). The x-axis represents years of projection and the y-axis represents total population; the grey band is the 10-90% confidence intervals, and the black line is the median of the population projection simulations. The black horizontal dotted line (***N***_f_) marks the population viability threshold used to estimate recovery potential.

### 5.3 Conservation management scenarios

We found partial support for our hypothesis H2: that recommended management scenarios would be inadequate for population recovery unless they increase adult survival by at least 80%. Our results indicate that management scenarios targeting either adults or juveniles in isolation are insufficient for population recovery, resulting in a population growth rate *λ*<1, which fails to shift the population trajectory upward toward the minimum viable population threshold (MVP; ***N***_f_ = 2,400), thereby failing to prevent functional extinction. However, (H2) is partially supported, as population recovery is achieved only under the combined management scenario (Scenario F: survival and fecundity both increased), contingent upon an 80% increase in adult survival (Fig. 3).

**Figure 3.**
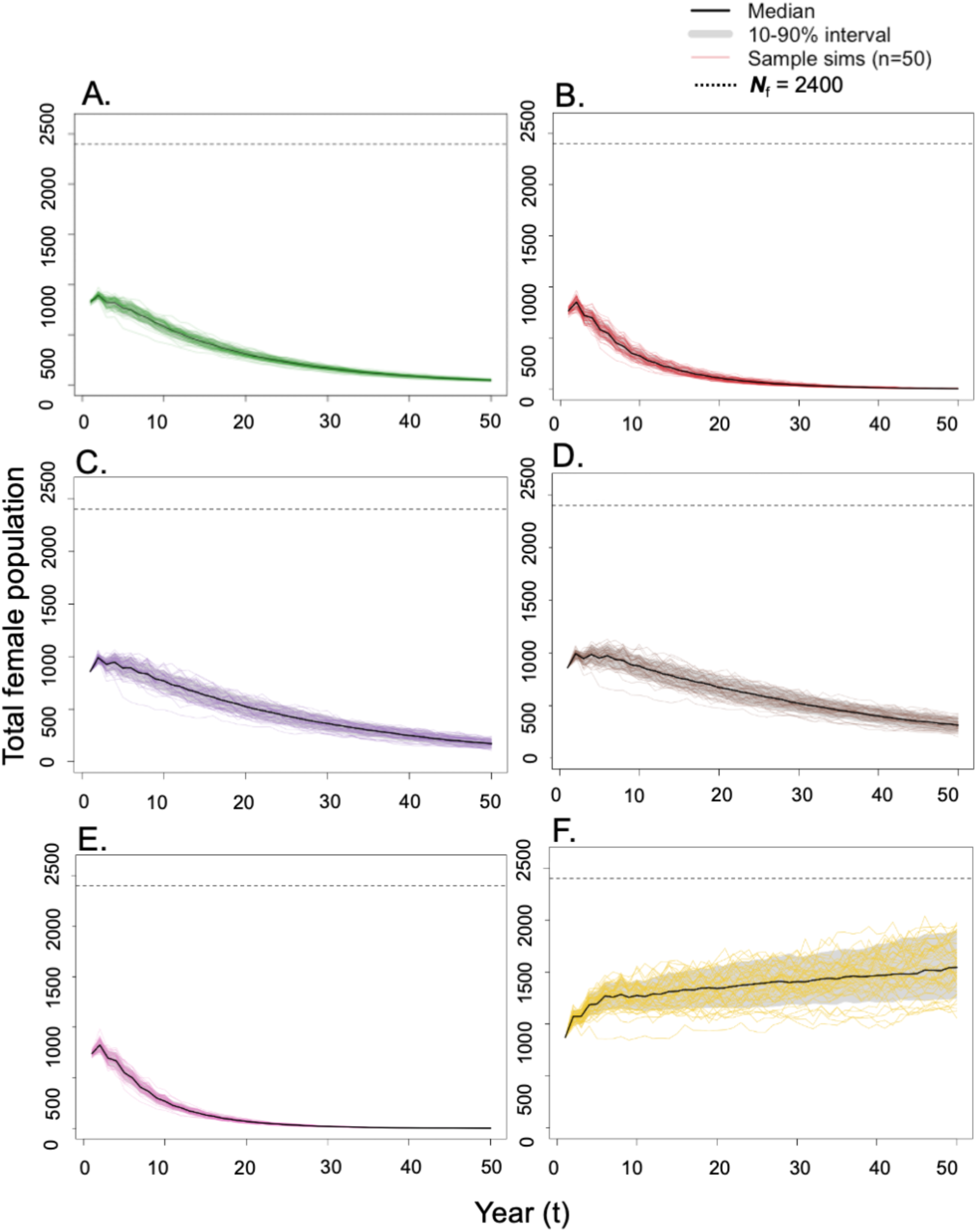
Population projections showing recovery potential under Scenario F for the state-structured Integral Projection Model for *Mobula mobular* population in India. Each scenario was stochastically projected for a 50-year time horizon (∼2 generations). Scenarios: **A.** Scenario A: Survival of Non-reproductive (NR) individuals is increased; **B.** Scenario B: Survival of Reproductive (R) and Sabbatical individuals is increased; **C.** Scenario C: Survival of all individuals is partially increased; **D.** Scenario D: Survival of all individuals is significantly increased; **E.** Scenario E: Fecundity of the population is partially increased; and **F.** Scenario F: Survival of all individuals and fecundity of the population is significantly increased. These population projections identify which scenarios avoid the risk of functional extinction and follow a trajectory toward recovery (>***N***_f_ : MVP threshold, black dotted line).

Scenarios A and B showed low effectiveness of policies targeting size-selective bycatch mitigation. We found a 6.9% increase in *λ*, resulting in a growth rate of 0.948 (0.938, 0.949 C.I.) (Table 3), following an 80% increase in Non-reproductive (NR) individuals under Scenario A. Projections over two generations (∼50 years) showed that, after an initial period of increase due to transient dynamics, the population declined gradually, reaching only a few hundred individuals within approximately one generation (Fig. 3). This suggests that despite a juvenile-dominant population structure, policy measures (Table 1) targeting juveniles alone, such as minimum size limits or market incentives deterring juvenile capture alone, are insufficient for population recovery or preventing functional extinction. Scenario B produced a lower *λ* than Scenario A, resulting in a *λ* of 0.917 (0.892, 0.918 C.I.) (Table 3), a 3.4% increase relative to the base model, approximately half the increase observed under Scenario A. Projections over 50 years showed a sharp decline under Scenario B, with the population trajectory declining sharply after approximately ten years (Fig. 3). Similar to the findings for Scenario A, this suggests that bycatch mitigation targeting only large individuals, such as gear compensation for releasing large individuals, maximum size limits for landings, and tRFMO safe-release requirements for large rays only (Table 1), is insufficient for recovery if applied in isolation.

Scenarios C and D showed moderately low effectiveness of policies targeting size-independent bycatch mitigation. Scenario C resulted in a higher *λ* when survival was partially increased across all size classes. Projections over a 50-year time horizon (approximately two generations) still indicated a decline; however, more gradual and resulting in larger population sizes compared to the size-targeted management scenarios (A and B) (Fig. 3). Scenario C resulted in a *λ* of 0.971 (0.959, 0.975 C.I.), a 9.5% increase relative to the base model (Table 3). This suggests that measures partially reducing bycatch (Table 1) such as the current partial protection of devil rays under schedule II & IV of the Wild Life (Protection) Act, 1972, mobulid retention quotas for domestic use, and incentives to release mobulid after catch quota is reached, are insufficient for recovery. Scenario D, applying a significantly increased survival across all three states (NR, R, and S), showed results similar to Scenario C, with a gradual decline (Fig. 3)—slower than Scenario C, as expected given the 80% survival increase as opposed to 50% in Scenario C. However, the population trajectory still did not shift upward toward the MVP (***N***_f_) threshold and continued to decline (*λ* = 0.981; 0.971, 0.985 C.I.). Scenario D showed the highest *λ* increase among all survival-targeting scenarios (A–D), a 10.6% increase relative to the base model (Table 3). Together scenarios A–D (Fig. 3), indicate that bycatch mitigation measures targeting survival alone (Table 1) are insufficient for population recovery unless complemented by measures targeting other vital processes in the *M. mobular* life cycle (Fig. 1).

We observed the lowest potential for population recovery under Scenario E, and the highest under Scenario F. When only fecundity was increased under Scenario E, *λ* showed the smallest increment relative to the base model, a 0.9% increase resulting in a *λ* of 0.895 (0.872, 0.893 C.I.) (Table 3; Fig. 3). This suggests that management strategies (Table 1) focused solely on increasing fecundity are insufficient for population recovery and are likely the least effective approach for recovering *M. mobular* populations when applied in isolation, without complementary measures improving survival. Scenario F, combining increased survival of individuals from all size classes with increased fecundity, is the only scenario showing recovery potential (Fig. 3). Over 50-years Scenario F demonstrates a shift in the population trajectory toward the MVP threshold (***N***_f_), thereby indicating the potential to prevent functional extinction over multiple generations. Under Scenario F, we found a 14.1% increase in population growth rate, with *λ* = 1.012 (1.004, 1.017 C.I.) (Table 3), with a gradual upward population trajectory (Fig. 3). Therefore, the upward population trajectory observed under Scenario F indicates that the only effective management strategy to recover populations of *M. mobular* is one that combines the highest levels of bycatch mitigation, resulting in significant increases in survival, with fisheries management measures that increase fecundity (Table 1).

### 5.4 Sensitivity analyses

We found partial support for H3: that the population growth rate (*λ*) for populations of *M. mobular* would exhibit high sensitivity to adult survival and that the population would have a heightened functional extinction risk when adult survival is low. Using parameter-level perturbations (Fig. 4), we found that *λ* shows moderately high sensitivity to survival of adults in the Reproductive (R) and Sabbatical (S) states. However, *λ* is most responsive to the slope of the reproduction vital rate function in reproductively active individuals (*φ*_1_), meaning *λ* is highly sensitive to changes in the relationship between individual size and reproductive output in females. Therefore, management measures affecting this relationship, for example those prioritising survival of large-sized reproducing females, would potentially have the largest effect on *λ*. However, *λ* also shows moderately high sensitivity to survival across all life stages (NR, R, S), including individuals in the NR state (Fig. 4). Therefore, long-term population recovery may only be possible if survival increases significantly across all size classes, especially for large reproducing females, with a concurrent increase in fecundity.

**Figure 4.**
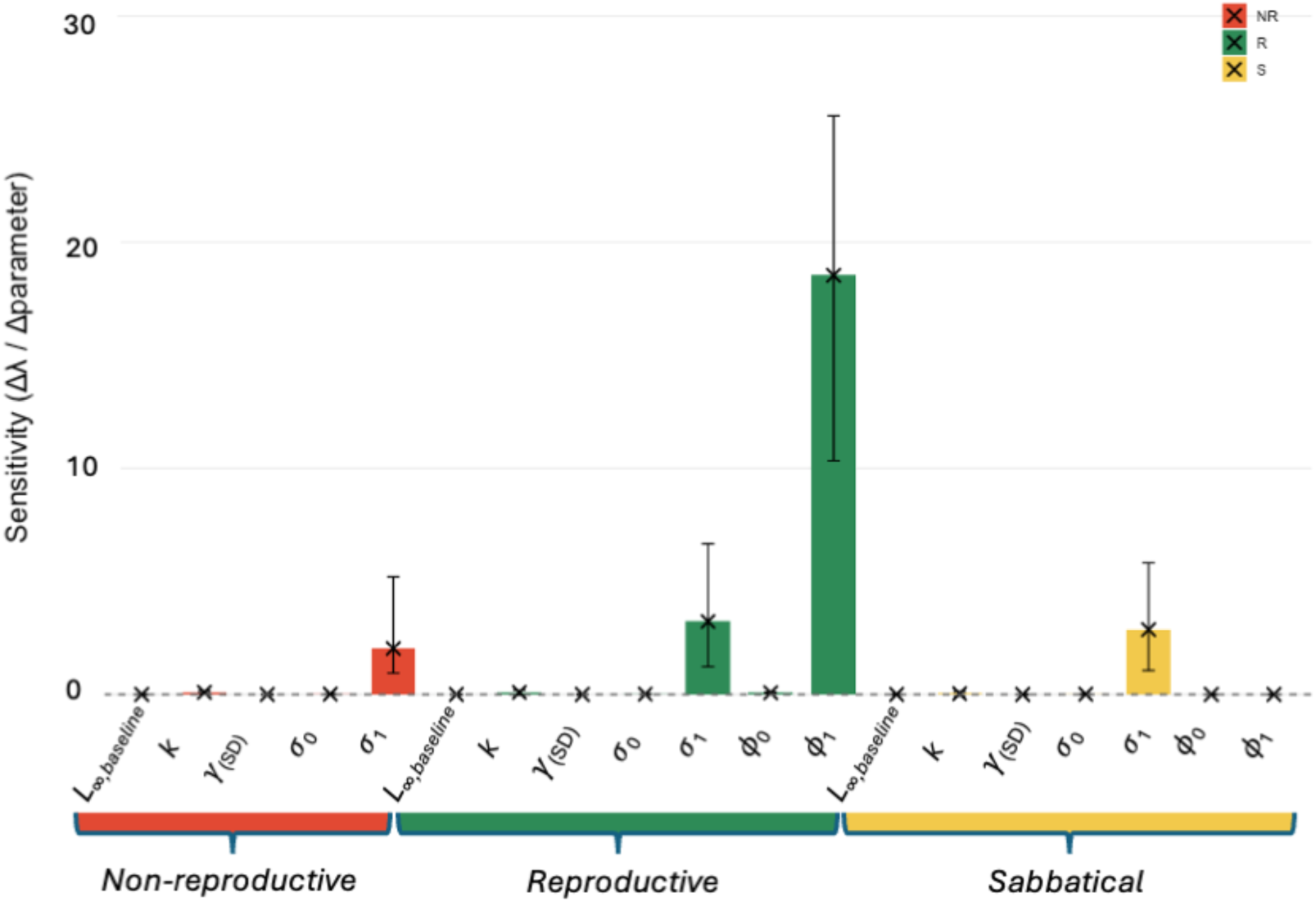
Parameter-level perturbations on the population growth rate (*λ*) show the highest sensitivity for the slope of the reproduction vital rate function in reproductively active individuals (*φ*_1_) and moderately high values for the slope (*σ*_1_) of the survival vital rate function across all three states: Non-reproductive (NR), Reproductive (R), and Sabbatical (S). The population growth rate (*λ*) estimates are for the female populations of *Mobula mobular* stochastic state-structured Integral Projection Model (IPM). The plot shows the sensitivity of parameter change to *λ* for each of the 19 parameters for survival, growth and fecundity, perturbed for all three life-cycle states (NR, R, S). The cross (×) represents the mean, and the error bars represent the 95% CI of sensitivity values for variations around baseline, using a simulation-based uncertainty propagation approach. The NR state vital rate parameters include, growth parameters (asymptotic growth rate [*L∞*_,baseline_]; Von Bertallanffy growth coefficient [*k*]; SD of growth transitions [*γ*_(*SD*)_]) and survival parameters (*σ*_0_, *σ*_1_). The R and S states additionally include two reproduction parameters (*φ*_0_, *φ*_1_).

## 6. Discussion

Human-induced disturbances have driven global biodiversity declines across taxa (Jungblut et al., 2020). Slow-living species have characteristically low resilience to disturbances (Capdevila et al., 2022; Dulvy et al., 2003; Lotze et al., 2011; Gamelon et al., 2014), so we assessed the risk of functional extinction and identified policy pathways for population recovery of the slow-living, Critically Endangered elasmobranch, the spinetail devil ray (*Mobula mobular*). Using a stochastic state-structured IPM (Easterling et al., 2000), we found no single policy measure sufficient to recover the *M. mobular* population along southeastern India’s coast; recovery requires integrated maximum bycatch mitigation and nursery area protection, targeting increased survival and fecundity together. The reported maximum intrinsic growth rate for Indian Ocean mobulid populations is *r_max_* = 0.109 yr^-1^ (an 11% annual increase under optimal conditions) (Barrowclift et al., 2025). In contrast, our base IPM estimated a finite population growth rate of log(*λ)* = −0.06 yr^-1^ (an approximate 12% annual decline) suggesting that the population is likely failing to approach its biological maximum growth potential due to overexploitation (Chopra et al., 2026). We found full support for (H1): *M. mobular* population will be at high-risk of functional extinction if current fishing pressure persists for another decade. We also found partial support for (H2): existing conservation measures are inadequate to facilitate population recovery unless they substantially increase adult survival for multiple generations, because our results show that both an increased adult and juvenile survival are required for recovery. Finally, we found partial support for (H3): *M. mobular* population exhibits high sensitivity to adult survival and have a heightened functional extinction risk when adult survival is low, as sensitivity to adult survival was high alongside high survival across all states and fecundity parameters.

The ‘business as usual’ harvest scenario, applying 86% harvest pressure over 10 years, resulted in large population declines. In support of H1, our results demonstrate a sharp decline leaving only a few hundred individuals after 10 years of sustained pressure. Notably, this decline occurred regardless of the variation in initial population size (***N***_0_ variation cases), showing limited influence of the mobulid population stock size on sustaining the population trajectory under continued harvest pressure. To prevent functional extinction and maintain evolutionary potential long-term, population census size (***N***_c_) needs to be in the thousands rather than hundreds (Brook et al., 2011; Clements et al., 2011; Traill et al., 2007), as population sizes of a few hundred can push the population into the so-called ‘extinction vortex’ (Gilpin and Soule, 1986; Brook et al., 2008). Therefore, we suggest that ‘business as usual’ harvest scenario should not continue, and that urgent management interventions are necessary to avoid local extirpation of these populations (Pardo et al., 2016).

Recovery from overexploited states often requires integrated resource management across life stages (Paterson et al., 2021; Keevil et al., 2023). In partial support of H2, our results identify Scenario F, as the only management action showing recovery potential. This may include, the inclusion and implementation of the highest national policy protection measure for mitigating bycatch of all mobulids, meaning protection under Schedule I of the Wild Life (Protection) Act, 1972 prohibiting all catch and trade (Bloch et al., 2016; Leduc and Hussey, 2019), alongside strict implementation of international protection translating to a complete international trade ban on mobulids and derived products under Appendix I of Convention of Trade in Endangered Species of Flora and Fauna (CITES) (O’Malley et al., 2017; Stewart et al., 2018; Palacios et al., 2024). The WLPA protection will also enable the implementation of existing obligations under CMS Appendices I and II, and the IOTC’s conservation and management measures (CMM) for mobulids. For example, the recent Concerted Actions for Manta and Devil rays adopted at the CMS CoP in 2026 (Convention on Migratory Species, 2026), call for CMS range states (including India) to work together on effective mobulid conservation management. We also recommend concurrent measures to improve fecundity, such as spatial protection of identified devil ray nursery areas and biologically relevant seasonal closures overlapping with mobulid reproductive periods (Gangal et al., 2023). Complementary measures enhancing juvenile and adult survival may further support recovery. For example, (i) market incentives for releasing caught juveniles, such as alternative meat sources discounts or socially-driven fisher incentives (Sun et al., 2017; Chopra et al., 2026); (ii) gear compensation for releasing large-sized mobulids to improve policy compliance (Bloch et al. 2016); (iii) training in safe handling and release using bycatch mitigation devices such as the manta grid (Cronin et al., 2024); and (iv) awareness campaigns for existing WLPA, 1972, CMS Appendices I & II, IOTC mobulid CMM, and CITES Appendix I protections. However, measures enhancing only juvenile or adult survival alone are unlikely to achieve long-term population recovery.

The need for integrative management, including bycatch mitigation and high-level nursery area protection, is further supported by the likelihood that this population is exhibiting “malediction,” typically of slow-living species (*sensu* Lebreton, 2006). Our forecasts also show transient dynamics, expected in the relative short-term following stochastic disturbance events (Stott et al., 2011; Gamelon et al., 2014), such as the overexploitation documented over the past decade (Chopra et al., 2026; Laglbauer et al., 2026). However, once these transient dynamics subside, population size continues to decline, consistent with demographic patterns reported for other slow-living species (Gamelon et al., 2014). Slow-living species are reportedly resilient to small magnitude disturbances, but large-scale negative events (Gamelon et al., 2014), such as declines of up to 99% in the last decade (CITES, 2025), likely decrease already low population growth rates and can push populations into the declining trajectory characteristic of the “extinction vortex” (*sensu* Lacy and Lindenmayer, 1995).

Population age structure typically has an important effect on growth rate and long-term viability (Wikström et al., 2012; Hoy et al., 2020). Changes to population structure following disturbances such as overexploitation and climate change have been widely linked to population declines across taxa (Kopf et al., 2025). The *M. mobular* population here is age-class truncated, with significant loss of larger individuals likely due to overfishing (Chopra et al., 2026). Given this right-skewed population structure, we show that although population growth rate is most sensitive to adult survival and reproduction parameters, increasing adult survival or reproduction alone does not shift the population trajectory towards recovery. This is likely because the population is predominantly juvenile, and applying protection only to the relatively small adult proportion is insufficient to drive long-term recovery (e.g., White et al., 2022). Consistent with low adult numbers, Scenario E (fecundity increase alone) shows the lowest long-term recovery potential, despite the reproduction parameter slope having the highest sensitivity to *λ*. We recommend measures that will enable maintaining a stable population age structure by increasing the survival of large-sized individuals, alongside protection across all size classes, as both are required for population persistence and long-term viability. Our recommendation aligns with context-specific management strategies proposed for other overexploited slow-living species (Wasser et al., 2017; Keevil et al., 2023), where recovery planning is structured around age-class contributions to population growth.

Equilibrium strategists, species characterised by slow growth, delayed maturation, and moderate to long lifespans, exhibit density-dependent vital rates (Stearns, 1976). They typically show large investment in offspring, including transfer of vital information from adults to juveniles (Nattrass et al., 2019) and the active regulation of social structures affecting fitness (Allen et al., 2021). In the absence of human harvest pressures, equilibrium-strategists’ life history traits have historically enabled their resilience to environmental variability (Pianka, 1970; Stearns, 1976). We argue that our results should be interpreted in the context of mobulids as equilibrium strategists, as they too are characterised by high per-offspring investment (one embryo per reproductive cycle; Villavicencio-Garayzar, 1991; Notarbartolo di Sciara, 1988) and slow maturation and growth (Barrowclift et al., 2025). For the *Mobula* genus, size is a good proxy for age (Rambahiniarison et al., 2018; Barrowclift et al., 2025), and thus size structure largely reflects age structure. As the life-history traits of equilibrium strategists are typically associated with stable age structures dominated by large and older individuals (Stearns, 1976), the size-class structure observed in our study system deviates from this expectation (similar to Fernando and Stewart, 2021). The deviation suggests that the population may no longer reflect its historical stable age distribution (Gamelon et al., 2014), potentially due to sustained harvesting pressure.

In slow-living equilibrium strategists, the population growth rate often shows high sensitivity to large adults (Stearns, 1976; Heppell et al., 2000). Our results partially support H3: population growth rate (*λ*) shows the highest sensitivity to changes in the relationship between individual size and reproductive output, and moderately high sensitivity to the relationship between individual size and survival of all size classes. Therefore, long-term population recovery for populations of *M. mobular* is only possible if survival increases significantly across all three life cycle states (Non-reproductive, Reproductive, Sabbatical) and size classes, especially for large reproducing females, with a concurrent increase in fecundity. Notably, we observed high sensitivities of *λ* to parameters denoting the relationship between individual size and survival (*σ*_1_), and reproduction (*φ*_1_). This higher importance of larger individuals is consistent with the general signature of sensitivities observed across slow-living wildlife (Cope et al., 2022). For example, in several slow-living mammal species, an abrupt fecundity decrease would shift *λ* from 1.1 to 1.0 causing an onset of increasing negative population momentum with increasing generation time (Gamelon et al. 2014).

Stock assessments are required to estimate initial population size and time to functional extinction (*e.g*., Davis, 2022). Here, we focused on recovery potential and management effectiveness based on the population structure derived from *M. mobular* bycatch data collected in 2023. We note that here we did not estimate the time to, or probability of, functional extinction, as these metrics may vary with total population size (Clements et al., 2011) and influenced by the connectivity among populations (Reed et al., 2002). The absence of a formal stock assessment (ICAR-CMFRI, 2022) means we lack an estimated total population size. Sparse information on connectivity between Indian Ocean mobulid populations further exacerbates this limitation for deriving time estimates to extinction (ICAR-CMFRI, 2022). Instead, we use the MVP (***N***_f_ = 2400) as an indicator for preventing functional extinction, identifying conservation management scenarios with recovery potential. We recommend that future work prioritise stock assessments for this population, enabling estimation of time to functional extinction and quantify the time available to implement the conservation measures proposed in this study.

In conclusion, the *M. mobular* population along southeastern India are exhibiting substantial annual declines, with overfishing inhibiting maximum growth potential. Current harvest pressures, if applied over the next 10 years, have a high-risk of pushing said populations into functional extinction (*sensu* Shaffer, 1981). Thus, “business as usual” harvesting should be replaced with urgent management interventions prioritising mobulid population recovery. Due to the existing age-class truncated population structure in the overexploited population stock, size targeted management measures applied in isolation do not show potential for population recovery. The examined *M. mobular* population shows long-term recovery (*λ>1*) only when maximum bycatch mitigation is complemented with spatial nursery area protection, calling for integrated and urgent action. We show that for slow-living species, loss of a stable age structure due to overexploitation likely makes recovery more challenging, with fewer viable scenarios. In the conservation context, a challenging population recovery means that higher management costs and efforts are required. Therefore, in data-limited settings, the precautionary principle, which emphasises preventive action under uncertainty, should guide conservation of slow-living species to prevent escalating management costs (Cooney and Dickson, 2005).

## 7. Acknowledgements

M.C. acknowledges the funding received through the Swami Vivekananda Scholarship for Academic Excellence (formerly Rajiv Gandhi Scholarship), granted by the State Government of Rajasthan, Government of India. Fieldwork and data collection were funded by a 2023 Graduate Student Research Award from the Society for Conservation Biology and the Emergency Grant provided by the Manta Trust. We thank our field assistants A. Nirojan, J. Dyson, and Ramesh in Tamil Nadu for their invaluable help with data collection and are especially grateful to the fishers at the landing centres of Thoothukudi, who cooperated when the mobulid landing data was being collected.

