## Supplementary Information 1 for "No single measure is enough: Recovery of the Critically Endangered *Mobula mobular* requires integrated maximum bycatch mitigation and nursery area protection"

### 1. Supplementary Information

### 1. Materials and methods supplementary

#### 1.1 Study site

Our study sites are located along the southeast coast of India (Figure 1A), in the Thoothukudi District, which has the second highest fishery landings in in the state (FRAEED-CMFRI, 2023), and records significant mobulid landings (Couturier et al., 2012; Sivadas et al., 2013). Data were collected at two sites within Thoothukudi District, Threspuram Fishing Village (8.8165° N, 78.1625° E) and Tharuvaikulam Fishing Harbour (8.8922° N, 78.1707° E) (Figure 1A), for six months between April and October in 2023. This mobulid bycatch (incidental catch in fisheries) fishery targets tuna and other pelagic species semi-selectively. Within this fishery, mobulids have exhibited a declining population trend over the past decade (Chopra et al., 2026). Sampling was designed to prioritise Thoothukudi's peak tuna fishing season (June–September), when mobulid landings are highest (Kumar, 2017). We monitored mobulid landings at jetties and fish landing centres during both morning and evening periods, averaging four hours of observation per day.

#### 1.2 Demographic data

We collected data from landed individuals in the study sites. For each individual, we recorded species, sex, and disc width. Disc width was measured as the distance between the tips of the pectoral fins without body curvature (Stevens et al., 2025). We did not assess female maturity because dissections were not performed, and assigned maturity information for our landed individuals using disc width maturity estimates from proximate mobulid populations (Rambahiniarison et al., 2018; D'Costa et al., in review). These data were recorded only when fishers provided explicit consent and when mobulid individuals were landed at fish landing centres instead of being transferred directly to transport vehicles for sale. When feasible, we photographed landed individuals, and three mobulid conservation experts independently crossverified species identification and demographic information using reference materials from Stevens et al. (2025).

#### 1.3 Vital rate estimation

To build the IPM, we first estimated parameters constituting the vital rates of survival, growth, and reproduction for the lifecycle of *M. mobular* (Figure 1).

***Survival***

To estimate state-specific baseline survival vital rates for the states, Non-reproductive (NR), Reproductive (R), and Sabbatical (S), we combined size-structured empirical data with literature-inferred survival estimates for *M. mobular*, and a closely-related species within the genus *Mobula*: *M. alfredi*. We inferred the yearling survival probability (0.63 $yr^{-1}$) from *M. alfredi*, which was estimated using capture-mark-recapture (CMR) analyses (Kashiwagi, 2014). As no disc width estimates were available for *M. mobular* yearlings (within the first year after birth), we adopted a conservative approach and assigned yearling survival probabilities to individuals with disc width <100 cm, corresponding to the minimum observed size at birth for *M. mobular* in an Indian Ocean population (Rambahiniarison et al., 2018). Similar to other elasmobranchs (Cortés, 2000), natural mortality in mobulids is higher during the first year of life, while juveniles and adults exhibit similar survival rates (Smallegange et al., 2016). As such, we used a survival probability of 0.93 $yr^{-1}$, representing the mean of published estimates for juvenile and adult *M. mobular* individuals (Pardo et al., 2016; Barrowclift et al., 2025), as an upper anchor point at adult size (217.8 cm). We used these literature-derived values to define a continuous logistic survival function of disc width, with annual survival probability $\sigma(z)$ expressed as:

$$\sigma(z) = \frac{1}{1+exp[-(\sigma_0+\sigma_1 z)]} \quad (1)$$

Where, $z$ denotes individual size and $\sigma_0$ and $\sigma_1$ are chosen survival parameters so that annual survival equals 0.63 at 100 cm and 0.93 at 217.8 cm. Using Eq. (1), we evaluated size-specific survival probabilities at size-bin midpoints and used these probabilities to simulate individual-level survival outcomes from Bernoulli trials. We then fitted a binomial generalised linear model to the simulated outcomes solely to obtain state-specific (NR, R, S) intercepts on the logit scale, while using a common size-dependent survival slope across the two states NR and R. The survival parameters of state S were assumed to be the same as state R because reproductively mature individuals alternatively transitioned between the two states (R and S).

***Growth***

For surviving individuals between times *t* and *t*+1, we next estimated growth parameters using the Von Bertalanffy Growth Function (VBGF). We used this approach as the VBGF provides the closest representation of the biological growth process for mobulids. Specifically, the VBGF represents growth that gradually slows down with increasing size, and eventually reaches an asymptotic maximum disc width (cm) (Dulvy et al., 2014b; Pardo et al., 2016; Rambahiniarison et al., 2018; Fernando & Stewart, 2021; Barrowclift et al., 2025). The asymptotic maximum disc width ($L\infty_{,\text{baseline}}$) is defined as the maximum size an individual would reach if it continued to grow indefinitely. As such, we set the asymptotic maximum disc width (cm), the annual growth coefficient, and the standard deviation of growth transitions (cm) around the expected disc width mean as per Eq. (2). The asymptotic maximum disc width ($L\infty_{,\text{baseline}}$) is estimated using Eq. (2) by Froese and Binohlan (2000):

$$L_{\infty,baseline} = 10^{0.044+0.9841\ log10\ (L_{max})} \qquad (2)$$

Where $L_{max}$ is the maximum observed disc width in 2023 for female populations of *M. mobular* (242 cm), and when solved for Eq. (2), provided an asymptotic maximum disc width of 245.4 cm. As we did not have paired observations of the same individuals from *t* and *t+1*, we could not have reliably estimated the growth coefficient (*k*) from our data. Therefore, we used the literature reported growth coefficient (*k* = 0.12) for populations of *M. mobular* (Pardo et al., 2016) in our VBGF. A standard deviation ($\gamma_{(SD)}$) of 6 cm was estimated around mean growth using the expected temporal variance through the growth function (Supplementary Information 1). Annual growth ($\gamma(z)$) to *t+1* for survivors was modelled as:

$$\gamma(z)\ = L_{\infty,baseline} - (L_{\infty,baseline} - z_t)\ exp(-kdt) \qquad (3)$$

Where $dt$ is 1 as the time interval for annual growth. As the asymptotic size and literature inferred growth rate are population level estimates, we set the same baseline growth parameters across all three states: NR, R, and S.

***Reproduction***

Females in the *Mobula* genus reproduce one embryo per reproductive cycle (Notarbartolo di Sciara, 1988; Villavicencio-Garayzar, 1991; Bradaï and Capapé, 2001; Guerrero-Maldonado, 2002; Serrano-López et al., 2021). Therefore, to model reproduction, we estimated vital rate parameters for reproduction probability ($\phi(z)$) of females in the NR state, the pup size ($D^{pup}$), and pup to recruitment probability ($p_{recruit}$). We defined reproduction probability ($\phi(z)$) as the probability of becoming pregnant for adult females in the NR state, and is expressed as:

$$\phi(z) = \frac{1}{1+exp[-(\phi_0+\phi_1 z)]} \tag{4}$$

Where $\phi_0$ and $\phi_1$ are reproduction probability parameters. To estimate newly recruited pup sizes, we assumed pup disc width at birth equals 50% of the maternal disc width, as inferred from the closely related species of *M. thurstoni* (Rambahiniarison et al., 2018). For each reproducing female, pup disc width at birth was:

$$D_{i,t}^{pup} = 0.5D_{i,t} \tag{5}$$

Where $D_i$ is the maternal disc width at time $t$. Recruitment probability ($p_{recruit}$) is the probability a newborn pup survives in the population to $t+1$. The pup to recruitment probability ($p_{recruit}$) of 0.6 was inferred from literature neonate survival from the closely related species of *M. alfredi* within the *Mobula* genus (Kashiwagi, 2014).

#### 1.4 Initialising the population vector

The $\boldsymbol{n_0}$ was derived from the 2023 landings data. As the observed size distribution in our 2023 data was right-skewed (with relatively few adults), we modelled the female disc width using a log-normal distribution. The parameters of this distribution were estimated from the log-transformed disc width data. The fitted distribution was then evaluated at the predefined IPM size-bin midpoints (200 mesh points per state) and scaled so that the values summed to one, giving the proportion of individuals in each size class. Initial counts in each size bin were then calculated as:

$$n_{0,i} = round(N_0 p_{z,i}), \tag{6}$$

With the first size bin adjusted to ensure that the total abundance summed exactly to $\boldsymbol{N_0}$, the initial population size.

#### 1.5 Estimating minimum viable population size threshold (MVP)

We considered population size approaching the MVP threshold ($\boldsymbol{N}_c$) of 5,000 individuals as an indication of the population being able to prevent the risk of becoming functionally extinct and show potential for recovery. To estimate the MVP for our modelled female population, we scaled this MVP ($\boldsymbol{N}_c$) to the observed proportion of females (0.48) in the populations of *M. mobular* in 2023, obtaining an MVP ($\boldsymbol{N}_f$) of 2,400 females.

#### 1.6 Estimation of growth variability ($\gamma_{(SD)}$)

To derive a biologically plausible estimate of growth variability ($\gamma_{(SD)}$) around the Von Bertalanffy Growth Function (VBGF), we calculated the expected one-year growth increment across the observed size distribution. We grouped observed disc widths into size bins, and the relative frequency of each bin was used to approximate the size structure of the population. For each size class midpoint *z*, the predicted disc width at the next time step was calculated using the VBGF, and the expected increment obtained was:

$$\Delta z = L_{t+1} - z, \tag{7}$$

A weighted mean increment was then computed using the observed size-class proportions, resulting in a mean predicted annual growth of 9.05 cm. We chose the $\gamma_{(SD)}$= 6cm to allow moderate inter-individual variability relative to the mean annual increment (9.05 cm), while maintaining biologically realistic growth estimates.

##### 1.7 Sensitivities to population growth rate

To quantify uncertainty around the sensitivity estimates, we implemented a simulation-based uncertainty propagation approach. Specifically, we simulated 200 stochastic parameter sets by varying each of the 19 baseline vital rate parameters around its baseline value using the same stochasticity values applied in the population projections. For each stochastic parameter set, we rebuilt the IPM and calculated $\lambda$. We then applied the same one-at-a-time perturbation ($\Delta\theta = 0.0001$) to each parameter in turn, maintaining all other parameters at their simulated values, and recalculated $\lambda$ to estimate sensitivities as per Eq. (5; stated in the main manuscript). This approach produced a distribution of sensitivity estimates for each parameter across 200 simulations. From these distributions, we calculated the mean sensitivity and 95% uncertainty intervals (Supplementary Table 2.1.1).

#### 2. Results supplementary

##### 2.1 Sensitivities to population growth rate

**Supplementary Table 2.1.1** High sensitivity for the slope of the reproduction vital rate function in reproductively active individuals ($\phi_1$), and moderately high values for the slope ($\sigma_1$) of the survival vital rate function across all three states (Non-reproductive (NR), Reproductive (R), Sabbatical (S)). Sensitivity values showing impact on population growth rate ($\lambda$) due to parameter level perturbations. The sensitivity values were estimated for the female *M. mobular* stochastic state-structured Integral Projection Model (IPM). The IPM is parameterised using demographic data collected in Tamil Nadu, India, and literature estimates for species within the *Mobula* genus. The table shows the sensitivity of parameter change to population growth rate ($\lambda$) when parameters for survival, growth and fecundity are perturbed for all life-cycle states (NR, R, S). The error bars represent the 95% CI of the sensitivity values for variations around the baseline, using a simulation-based uncertainty propagation approach. The vital rate parameters of the Non-reproductive (NR) state include, growth parameters: asymptotic growth rate ($L\infty_{,baseline}$), Von Bertallanffy growth coefficient ($k$), SD of growth transitions ($\gamma_{(SD)}$); and survival parameters ($\sigma_0$, $\sigma_1$). In addition to the growth and survival parameters, the Reproductive (R) and Sabbatical (S) states also have two reproduction parameters ($\phi_0$, $\phi_1$).

| **Vital rate parameter** | **Baseline Value** | **Baseline sensitivity** | **Sensitivity mean** | **Sensitivity (lower limit CI)** | **Sensitivity (upper limit CI)** | **Life-cycle state** |
|---|---|---|---|---|---|---|
| Survival ($\sigma_0$) | -1.84 | 0.01155 | 0.01402 | 0.00650 | 0.03516 | Non-reproductive |
| Survival ($\sigma_1$) | 0.021 | 1.66607 | 2.03571 | 0.93507 | 5.21035 | Non-reproductive |
| Asymptotic size ($L_{\infty,baseline}$) | 245.4 cm | 0.00011 | 0.00012 | 0.00006 | 0.00024 | Non-reproductive |
| Growth coefficient *(k)* | 0.12 | 0.08689 | 0.09234 | 0.05168 | 0.15529 | Non-reproductive |
| Standard deviation $\gamma_{(SD)}$ | 6 cm | 0.00006 | 0.00005 | 0.00001 | 0.00010 | Non-reproductive |
| Survival ($\sigma_0$) | -1.63 | 0.01354 | 0.01416 | 0.00560 | 0.02932 | Reproductive |
| Survival ($\sigma_1$) | 0.021 | 3.09033 | 3.23366 | 1.22520 | 6.66477 | Reproductive |
| Asymptotic size ($L_{\infty,baseline}$) | 245.4 cm | 0.00121 | 0.00119 | 0.00084 | 0.00154 | Reproductive |
| Growth coefficient *(k)* | 0.12 | 0.08884 | 0.09114 | -0.03020 | 0.19061 | Reproductive |
| Standard deviation $\gamma_{(SD)}$ | 6 cm | 0.00035 | 0.00043 | -0.00090 | 0.00198 | Reproductive |

| | | | | | | |
|---|---|---|---|---|---|---|
| Reproduction ($\phi_0$) | -6 | 0.07515 | 0.07381 | 0.04284 | 0.10161 | Reproductive |
| Reproduction ($\phi_1$) | 0.03 | 18.62826 | 18.51699 | 10.33520 | 25.59154 | Reproductive |
| Survival ($\sigma_0$) | -1.63 | 0.01193 | 0.01215 | 0.00440 | 0.02491 | Sabbatical |
| Survival ($\sigma_1$) | 0.021 | 2.82649 | 2.87628 | 1.05335 | 5.82658 | Sabbatical |
| Asymptotic size ($L_{\infty,\text{baseline}}$) | 245.4 cm | 0.00127 | 0.00123 | 0.00089 | 0.00158 | Sabbatical |
| Growth coefficient *(k)* | 0.12 | 0.03869 | 0.03439 | -0.07160 | 0.12761 | Sabbatical |
| Standard deviation $\gamma_{(SD)}$ | 6 cm | 0.00035 | 0.00044 | -0.00107 | 0.00224 | Sabbatical |
| Reproduction ($\phi_0$) | 0.03 | 0 | 0 | 0 | 0 | Sabbatical |
| Reproduction ($\phi_1$) | 0.03 | 0 | 0 | 0 | 0 | Sabbatical |
